# Miniaturized CAR knocked onto CD3ε extends TCR function with CAR specificity under control of endogenous TCR signaling cascade

**DOI:** 10.1101/2023.05.30.542829

**Authors:** Katrin Manske, Lisa Dressler, Simon P. Fräßle, Manuel Effenberger, Claudia Tschulik, Vlad Cletiu, Eileen Benke, Michaela Wagner, Kilian Schober, Thomas R. Müller, Dirk H. Busch, Christian Stemberger, Lothar Germeroth, Mateusz P. Poltorak

## Abstract

Immunotherapy using TCR and especially CAR transgenic T cells is a rapidly advancing field with the potential to become standard of care for the treatment of multiple diseases. While all current FDA approved CAR T cell products are generated using lentiviral gene transfer, extensive work is put into CRISPR/Cas mediated gene delivery to develop the next generation of safer and more potent cell products. One limitation of all editing systems is the size restriction of the knock-in cargo. Targeted integration under control of an endogenous promotor and/or signaling cascades opens the possibility to reduce CAR gene size to absolute minimum. Here we demonstrate that a first-generation CAR payload can be reduced to its minimum component - the antigen-binding domain - by targeted integration under control of the CD3ε promoter generating a CAR-CD3ε fusion protein that exploits the endogenous TCR signaling cascade. Miniaturizing CAR payload in this way results in potent CAR activity while simultaneously retaining the primary antigen recognition function of the TCR. Introducing CAR-specificity using a CAR binder only while maintaining endogenous TCR function may be an appealing design for future autologous CAR T cell therapies.

## Introduction

T cells engineered to express engineered T cell receptor (eTCR) or chimeric antigen receptor (CAR) constructs have become a powerful tool in the field of immunotherapy. As of today, multiple T cell therapies are approved by the Food and Drug Administration (FDA) and European Medicines Agency (EMA) showing impressive therapeutic effect and curative potential for patients with different types of blood cancers ^1,2,3,4,5,6^. While it’s the midterm goal to make CAR T cell therapy more accessible by lowering manufacturing cost and to broaden its applicability to new tumor entities including solid tumors, the field is also refining techniques to engineer CAR T cells to produce safer and more potent cell products. Clustered Regularly Interspaced Short Palindromic Repeats (CRISPR)/Cas may pave the way to this sophisticated and safer gene engineering and holds the promise to revolutionize future cell manufacturing.

More advanced engineered cell therapies employ either CARs with multiple targeting domains, additional genes to overcome the immunosuppressive tumor microenvironment in solid tumors, or genes designed to mitigate host rejection of allogeneic cell therapies ^6,7^. These payloads are considerably larger than CARs employed in approved therapies and are typically delivered to targeted genomic loci using CRISPR/Cas-mediated editing via either adeno-associated virus (AAV) or dsDNA/ssDNA non-viral delivery templates. The packaging limit of AAV is ∼5kb and non-viral ssDNA and dsDNA templates introduce higher toxicity and lower efficiency as payload size increases. Although lentivirus (LV) has a relatively large payload capacity (up to 9kb), it is not compatible with targeted delivery to a specific locus. Techniques that deliver similar performance with a smaller genomic payload could therefore improve both the potency and manufacturability of these more advanced engineered cell therapies. Careful selection of the targeted integration site may allow significantly reduction of overall construct size if it is placed under control of endogenous promoter and/or downstream signaling domains.

An intuitive target sequence for either eTCR or CAR constructs are TCR/CD3 complex-associated loci. The TCR/CD3 complex is one of the most intricate receptor structures 8. Each receptor complex consists of six different subunits with TCRα and TCRβ chains being responsible for antigen recognition via peptide MHC (pMHC) binding 9,10. The TCRαβ subunits are associated with CD3 hetero or homodimers formed by CD3εγ, CD3εδ, and CD3ζζ to initiate the downstream T cell signaling cascade 11. It has been shown that targeting a CAR or eTCR construct into the endogenous TCR locus optimizes their expression, safety, and function ^12^. Additionally, it was demonstrated that fusion molecules combining CARs scFv fragment and TCR/CD3 complex can replace full-length CARs ^13,14^. Similarly, the eTCR or CAR sequence can be minimized to the antigen-recognition site only (either TRAV and TRBV for eTCR or VH and VL scFv sequence for CAR, respectively), if it is placed into TCR locus itself to take advantage of endogenous intracellular TCR signaling pathways ^15,16^.

Here, we demonstrate that the anti-CD19 scFv of a CAR construct can be integrated into the CD3ε locus generating a CAR-CD3ε fusion protein which preserves the native TCR structure. Moreover, by using the endogenous promoter and signaling cascade we reduce the genetic payload of the CAR to its minimum, the antigen-binding site. In this manuscript, we present the proof-of-concept data for this minimal CAR construct (miniCAR) and evaluate its benefits in both *in vitro* and *in vivo* studies.

## Results

The present study demonstrates a method to exploit the endogenous TCR/CD3 complex to co-express a scFv fragment of a defined specificity using CRISPR/Cas and targeted integration via non-viral gene delivery.

In classical LV transduction approaches, a full-length CAR construct including an external promoter (LV CAR) is randomly integrated in the host cell (Fig 1A, upper schematic). Newer approaches introduce the CAR constructs into the TRAC locus using CRISPR/Cas homology directed repair (HDR)-mediated targeted integration system. Here, CAR expression is driven by the endogenous TRAC promoter thus the integrated gene is devoid of an external promoter but still requires the complete artificial signaling cascades and polyA tail of classical LV CAR constructs and destroys TRAC gene codon sequence preventing TCR/CD3 assembly on the cell surface (Fig 1A, middle schematic). In contrast, our approach specifically targets the CAR to the CD3ε locus 3’
s of the native promotor region (Figure 1A, lower schematic). The HDR template is designed to fuse the CAR scFv including a small linker seamlessly upstream to the fully preserved CD3ε (Fig 1A). This concept allows us to reduce the first-generation CAR construct to its minimum: the scFV fragment (henceforth named miniCAR) and the signaling domains of CD3ε are used to promote cell activation via the endogenous TCR signaling cascade (Figure 1A, lower row) ^15,16^.

**Figure 1:**
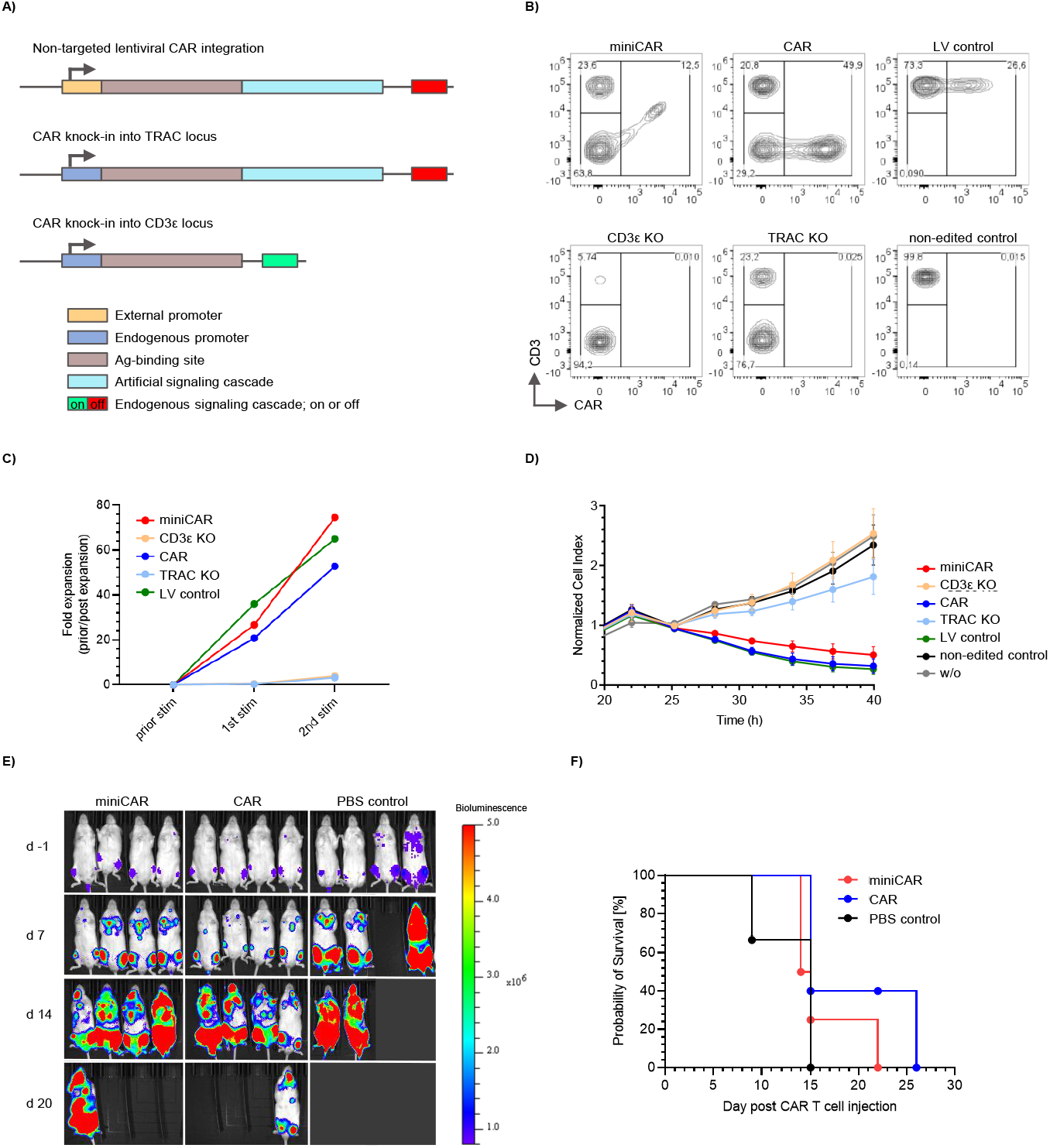
Miniaturized CAR (miniCAR) exploits endogenous signaling cascade generating fully functional CAR T cells. **A)** Schematic illustration of anti-CD19-CAR constructs. Upper line: non-targeted CAR sequence integration into the genome after lentiviral transduction (LV control). The CAR expression is regulated by an exogenous promoter. Middle line: non-viral CRISPR-mediated CAR knock-in into the TRAC locus resulting in T cell receptor knock-out (CAR). Lower line: non-viral CRISPR-mediated miniaturized CAR-CD3ε knock-in into CD3ε locus reassembling the CD3 sequence and exploiting endogenous T cell receptor signaling (miniCAR). **B)** Flow-cytometric analysis of CD3 and CAR expression on primary human T cells on day 12 post gene-editing by CRISPR technology or lentiviral transduction. **C)** Fold expansion of primary human CAR T cells post antigen-dependent stimulation. For anti-CD19-CAR stimulation, T cells were co-cultured with LCLs at a E:T ratio of 1:7. **D)** Cytotoxicity of primary human CAR T cells during co-culture with CD19-expressing HEK293 tumor cells measured by xCELLigence RTCA System. HEK293 cells and T cells were co-cultured at E:T ratio of 2.5:1. **E)** Bioluminescence imaging of tumor-bearing NSG-SGM3 mice treated by human miniCAR T cells, CAR T cells or PBS as control. Mice were engrafted with 5e5 CD19+ Raji/ffluc-GFP tumor cells at day -1. At day 7, the mice were treated by 1e6 miniCAR or CAR T cells. The scale indicates bioluminescence signal at the range of 8e5-5e6 p/s/cm2/sr. **E)** Survival probability of tumor-bearing NSG-SGM3 mice treated as in E).

For our study we used a publicly available anti-CD19 scFv fragment which is widely used as a binder. This enabled us to evaluate the miniCAR in well-established experimental settings. First, we assessed the expression of the miniCAR on the surface of primary human T cells with flow cytometry (Figure 1B). We could efficiently knock out CD3ε (Figure 1B, lower left panel). When combining the ribonucleoprotein (RNP) directed against CD3E locus with HDR template encoding the miniCAR sequence we detected double positive staining for CAR and CD3 indicating expression of a miniCAR-CD3ε fusion protein (Figure 1B, upper left panel). In contrast, TRAC knock-out resulted in loss of CD3 on the surface (Figure 1B, lower middle panel), which was not restored upon CAR KI into TRAC locus (Figure 1B upper middle panel). As expected, LV transduction did not alter CD3 expression (Figure 1B right panels). These data shows that knock-in miniCAR construct into the CD3E locus successfully generates the surface expression of a CAR-CD3ε molecule on primary human T cells.

Next, we compared the CD19 antigen recognition and subsequent proliferative potential of primary human T cells expressing either miniCAR knocked-onto the CD3ε or first-generation CAR knocked-in the TRAC locus, respectively (Figure 1C). Both modified cell populations reacted to antigen-specific stimulation (CD19^+^ LCL cell line) comparable to LV control. We further tested the cytolytic function of miniCAR T cells directed against CD19 expressing target cells (Figure 1D). In an *in vitro* killing assay miniCAR T cells were as efficient in target cell lysis as their full-length CAR T cell counterparts in an *in vitro* killing assay (Figure 1D). Based on the encouraging results, showing comparable proliferative potential and strong *in vitro* killing capacity, we investigated the *in vivo* potential of miniCAR T cells in comparison to classical CAR T cells. To better tease out potential differences between these groups, mice were injected with a sub-curative dose of CAR^+^ T cells. After injection mice were imaged and blood was analyzed once a week. Compared to PBS control, mice injected with miniCAR as well as full-length CAR T cells showed a survival benefit but were not able to control or fully clear the tumor (Figure 1E). However, the number of transferred T cells and CAR^+^ T cells in the blood increased over time, indicating an antigen specific response (Supplementary Figure 1). These results demonstrate functionality of miniCAR when compared to full-length CAR with minimal advantage of the latter.

Next, we aimed to investigate whether miniCAR T cells preserve the endogenous TCR/CD3 complex. To this end, we first performed a more detailed analysis of miniCAR transgenic primary human T cells (Figure 2A). As anticipated, knock-out of CD3ε led to complete depletion of TCR/CD3 complexes (Figure 1B, bottom left panel). Importantly, miniCAR knock-in partially rescued not only CD3, but led to increased TCRαβ detection as well (albeit not to the previous intensity) (Figure 1B and Figure 2A). Lower TCRαβ signal intensity can hint toward reduced expression rates of the miniCAR-CD3ε fusion protein, suboptimal TCR complex formation especially in polyclonal T cell populations, or a technical artifact caused by antigen competition/masking as a result of steric hindrance of detection antibodies in high proximity. Nevertheless, we can speculate that the observed TCRαβ expression range indicates a potentially preserved cognate antigen binding function of restored TCR complexes.

**Figure 2:**
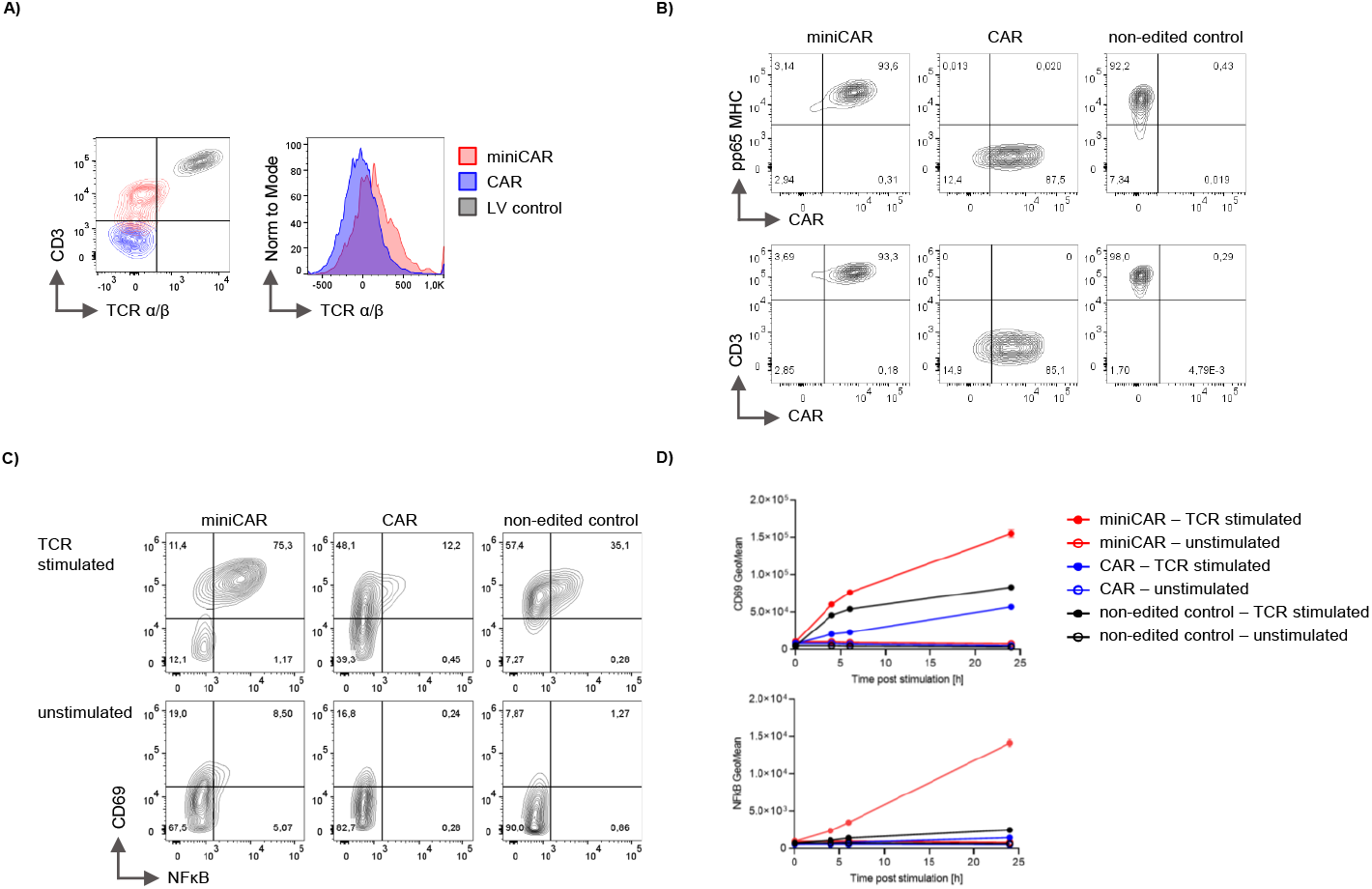
Reconstitution of TCR-mediated signaling in engineered miniCAR T cells. **A)** Flow-cytometric analysis of CD3 and TCRαβ expression on primary human T cells on day 12 post gene-editing. **B)** Flow-cytometric analysis of NFκB-reporter Jurkats expressing miniCAR or CAR introduced by CRISPR-mediated gene-editing. CAR, TCR and CD3 expression were analyzed respectively. CMV-specific TCR was detected by A2/pp65 pMHC multimer. **C)** NFκB-reporter Jurkats harboring the CMV-specific TCR were edited with miniCAR or CAR by CRISPR-technology. CD69 and NFκB expression were detected by flow-cytometry post stimulation with A2/pp65 pMHC multimer and anti-CD28-Fabs. Flow-cytometric plots show the CD69 and NFκB expression 6h post stimulation. Geometric Mean of CD69 and NFκB expression was quantified at 4h, 6h and 24h post stimulation.

To investigate the functionality of TCR/CD3 complexes carrying miniCAR in a better controlled system, we used a modified Jurkat T cell line expressing an engineered TCR of known specificity (A2/pp65) introduced into TRAC locus as well as NFκB reporter (Figure 2B-D). First, we assessed pMHC multimer loaded with cognate peptide binding before and after CAR editing (Figure 2B, upper panels). miniCAR transgenic T cells were able to bind A2/pp65 pMHC multimer similarly to the non-edited control cells indicating unaltered antigen recognition via the TCR/CD3 complex carrying miniCAR. Conversely, CAR transgenic T cells were unable to bind A2/pp65 pMHC multimer since the gene is knocked into the TRAC locus (Figure 2B, upper panels). Thus, in the monoclonal TCR setting, miniCAR expression directly correlates with cognate antigen recognition via the TCR complex (Figure 2B, lower panel).

Next, we investigated whether the detectable TCR/CD3 complex and maintained pMHC multimer recognition translates into TCR-mediated responses. Hence, we stimulated the CAR transgenic cells lines with TCR-specific stimulus (Figure 2C). Interestingly, both miniCAR edited, and non-edited cells responded to TCR-restricted stimulus as indicated by upregulation of CD69 and NFκB (Figure 1C, upper left and right panel). Full-length CAR control samples did show background level activation, which might be explained by antigen independent anti-CD28 stimulus or residual/cycling TCR^+^ Jurkats in the purified cell line population (Figure 2C, middle panel). These data confirmed maintenance/restoration of endogenous TCR capacity after integration of miniCAR into the TCR/CD3 complex in mono and polyclonal settings.

In summary, we demonstrate through *in vitro* and *in vivo* assays that first-generation CAR constructs can be reduced to its absolute minimum - the scFv fragment - when targeting it under the control of endogenous promotor and signaling cascade of CD3ε.

## Discussion

Here, we describe that by careful selection of the integration site transgenes can be reduced to their absolute minimum when exploiting endogenous promoters and signaling domains. Moreover, the manipulated native gene/protein remains fully intact resulting in gain-of-function in the transgenic cell. For proof-of-concept experiments we focused on a clinically relevant artificial gene for T cell modification in immunotherapy – the Chimeric Antigen Receptor (CAR). In this setting, the sole scFV fragment of a CAR was placed under control of the CD3ε promoter generating a CAR-CD3ε fusion protein. Consequently, the CAR became an integral part of the TCR/CD3 complex and used the endogenous TCR signaling cascade for activation signal transmission. This approach not only opens a way to more physiological signal transduction for CAR T cell engineering but reduces the necessary genetic payload. The novelty of our approach is the means of creating the fusion protein on a cellular level and not to modify cells with the fusion protein itself.

Smaller transgenes can be of great interest when considering the size restrictions of certain gene transfer technologies. The emergence of dual-antigen specific CAR constructs and additional modifications to armor transgenic T cells to overcome the tumor suppressive microenvironment increase the need to reduce genetic cargo by clever integration and construct design ^17,18,19^. Transgene size and efficiency of gene-transfer correlates for most transfer systems and emerging DNA/RNA vehicles such as AAVs and LNPs have relatively low cargo load capacity in general ^20,21,22^. Same accounts for the length of HDR templates used for virus-free CRISPR/Cas gene transfer via electroporation ^23^.

We demonstrate that the miniCAR constructs can be efficiently expressed on cell surface of engineered T cells. When co-cultured with CD19^+^ target cells *in vitro*, miniCAR T cells respond with strong antigen-depended T cell proliferation as well as control of target growth. The miniCAR *in vitro* function was highly comparable to their full-length first-generation CAR construct counterpart. Importantly, we could also demonstrate that miniCAR T cells provide survival benefit in murine Raji tumor model comparable to full-length CAR KI T cells even when applying a sub-curative dose level. These results demonstrated that the introduced miniCAR construct exerts full functionality although being more than 50% reduced in size. It has been shown that native-like regulated TCR/CAR transgenic T cells are more potent than CAR T cells with constitutive active CAR expression. Although we were not able to detect any substantial advantage of miniCAR, it is reasonable to speculate that exploiting the endogenous signaling cascade by an introduced miniCAR construct can lead to long-term benefits as previously shown in in vivo models ^15^.

Our proof-of-concept data demonstrate the feasibility of placing an artificial gene under the control of endogenous promotor and signaling domains. This design might allow getting one step closer to a native-like integration of exogenous genes into physiological signaling cascades. It has been shown that native-like regulated TCR/CAR transgenic T cells are more potent than CAR T cells with constitutive active CAR expression ^24,25^. Another advantage of using an endogenous promoter to drive transgene expression is safety. Choosing a cell type specific promotor, such as TCR related genes for T cell or B-cell receptor related genes for B cells, reduces the risk of gene expression in off-target cells ^25^.

In contrast to most CRISPR/Cas-mediated CAR engineering approaches our modification strategy does not aim to knock-out endogenous genes. This would in theory allow transgenic cells to maintain some native functionality while gaining additional function mediated by the gene-of-interest. This idea is supported by the finding, we were able to show that the miniCAR-CD3ε fusion protein allows partial rescue of the native TCR complex in polyclonal primary human T cells as well as monoclonal Jurkat cell line with a defined TCR. Interestingly, miniCAR transgenic Jurkat cells were able to bind their cognate pMHC multimer as efficiently as non-edited Jurkats indicating that the antigen-recognition is fully preserved which was underlined by antigen-specific upregulation of T cell activation markers CD69 and NFκB. Even if our data illustrates that miniCAR T cells modified T cells are bi-functional, we cannot exclude the possibility that dual-antigen specificity might be restricted to particular TCRs. A follow up evaluation with multiple pre-defined TCR complexes will be needed to fully address this statement.

In conclusion, the proposed miniCAR design points to a strategy to embed artificial gene expression and function under the control of endogenous genes. We believe that the described manipulation strategy can help to generate better cell products and facilitate more efficient manufacturing through reduced transgene size.

## Materials & Methods

### Human blood donors

T cells were isolated from leukapheresis collected from healthy donors by CCC Cellex Collection Center Dresden considering ethical quote (Ethical committee of the Technical University Dresden: EK309072016). Donor samples were approved by national law by the local Institutional Review Board and the declaration of Helsinki and Istanbul (Ethics committee of the Faculty of Medicine, Technical University of Munich: 360/13 and 55/14).

### T cell selection and activation

Leukapheresis was washed with 1x PBS (Thermo Fischer Scientific) containing 0.5% HSA (CSL Behring) by automated cell separation system Sepax S-100 (Cytiva). Further, T cells were selected and activated for 4.5 h by the in-house T cell selection technology ATC as described in Radish and Poltorak et al. ^26^. Selected T cells were cultured in serum-free media overnight in presence of 100 IU/ml recombinant human IL-2, 600 IU/ml recombinant human IL-7, 100 IU/ml recombinant human IL-15 (R&D Systems) and aCD3/aCD28 Expamers as previously described ^27^.

### T cell phenotyping

T cells were analyzed by flow-cytometry using the anti-CD45 (PE; clone HI300), anti-CD3 (PE, FITC; clone OKT3), anti-CD8 (BV510; clone HIT8a), anti-CD4 (BV421; clone OKT4), anti-CD69 (APC-Fire750; clone FN50) and anti-CD25 (BV650; clone BC96) (Biolegend) and anti-idiotype antibody detecting anti-CD19-CAR coupled to AF647 (Bristol-Myers Squibb). Dead cells were discriminated by propidium iodide (Thermo Fisher Scientific). Flow-cytometric analysis was performed with CytoFLEX or CytoFLEX S flow-cytometer (Beckman Coulter).

### Cell culture

Primary human T cells were cultured in serum-free media (Thermo Fisher) containing 100 IU/ml recombinant human IL-2, 600 IU/ml recombinant human IL-7, 100 IU/ml recombinant human IL-15 (R&D Systems). NFκB-reporter Jurkat cell line expressing cyan fluorescent protein (CFP) under control of NFκB promoter was cultured in RPMI 1640 (Gibco), 10% heat-inactivated FBS (Gibco), 1% P/S (Gibco), 1% glutamine (Gibco), 1% NEAA (Gibco). Human Embryonic Kidney cells (HEK293-CD19+) were cultured in DMEM (Gibco), 10% heat-inactivated FBS (Gibco), 1% P/S (Gibco), 1% glutamine (Gibco), 1% NEAA (Gibco).

### Gene editing

T cells were gene-edited by CAR knock-in into the TRAC locus resulting in TCR disruption. In contrast, miniCAR was knocked-in into the CD3ε locus reassembling the CD3 protein and maintaining the TCR expression. T cells (24 h activated) and Jurkats, were electroporated in presence of 1 μg double-stranded linear DNA template and 3 μl Cas9 ribonucleoproteins per 1e6 cells. Electroporation of primary T cells was performed in Nucleofector Solution P3 with 4D Nucleofector X unit, program code EH-100 (Lonza). Post electroporation, primary T cells were rested in cytokine free T cell medium 15 min at 37°C. Subsequently, primary T cells were cultured overnight in presence of 100 IU/ml recombinant human IL-2, 600 IU/ml recombinant human IL-7, 100 IU/ml recombinant human IL-15 (R&D Systems) and small molecule inhibitors 1 μM XL-413 and M-3814 (Selleck Chemicals). 16 h post culture, small molecule inhibitors were removed by medium exchange. Electroporation of Jurkats was performed in SE Cell Line Nucleofector Solution with 4D Nucleofector X unit, program code CL-120 (Lonza) and cultured in RPMI medium without small molecule inhibitors. CAR-expressing Jurkat population was enriched by bulk cell sort using the CytoFLEX SRT (Beckman Coulter).

For non-targeted anti-CD19-CAR integration, activated human primary cells were transduced with replication-deficient lentiviral particles encoding anti-CD19-CAR in cell culture (Bristol-Myers Squibb Company’s third-party supplier).

### RNP generation

For generation of gRNA, 80 μM crRNA (IDT DNA) and 80 μM crRNA (IDT DNA) were incubated 5 min at 95°C. After cooling down to room temperature, high fidelity Cas9 (IDT DNA) at a final concentration of 3 μM and electroporation enhancer (IDT DNA) at a final concentration of 30 μM were added to the gRNA mix. The RNPs were assembled 20 min at room temperature and used directly for gene-editing.

### DNA template generation

Double-stranded linear DNA template for HDR and electroporation was amplified from plasmid DNA by PCR using Q5 High-Fidelity Master Mix (New England Biolabs) according to manufacturer’s protocol. We used two sets of primers for either CAR or miniCAR DNA double-stranded linear DNA templates. Both primer pairs were used at were used at a final concentration of 0.5 μM. For generation of DNA double-stranded linear DNA template, thermal cycling was performed as follows: denaturation at 98°C 30 sec, 36x cycles of denaturation (98°C 10 min), annealing (60°C 20 s), extension (72°C 2 min) and final extension at 72°C 3 min. The PCR product was purified by Ampure XP beads (Beckman Coulter) according to manufacturer’s protocol.

### Cytotoxicity assay

20,000 Human Embryonic Kidney cells (HEK293-CD19+) expressing the CD19 antigen were seeded on a 96-well E-plate (ACEA Biosciences) over night. On the next day, rested human primary gene-edited T cells were added at a ratio of 5:1 (E:T). Target cell death was measured by impedance-based xCELLigence RTCA System (ACEA Biosciences).

### Cell stimulation assay

For cell stimulation, Jurkats were cultured at a density of 1e6/ml in RPMI medium in presence of 4 μg CMV-specific pMHC/anti-CD28 as previously described ^28^. Cells were stimulated for 4, 6 and 24 h. Cell activation was detected by flow-cytometry measuring CD69 and NFκB expression. NFκB expression was assessed by the reporter gene cyan fluorescent protein.

For characterization of expansion, gene-edited primary human T cells were co-cultured with irradiated CD19^+^ lymphoblastoid cell line (LCL) in a ratio of 1:7 (E:T) as described in Turtle et al.,^29^.

### Mice

NSG-SGM3 (NOD.Cg-Prkdcscid Il2rgtmWjlTg (CMV-IL3, CSF2, KITLG) 1Eav/MloySzJ, NSGS) mice were purchased from Jackson Laboratory and maintained in the animal facility of Technical University Munich under specific pathogen-free conditions. Guidelines of the Federation of Laboratory Animal Science Association were applied, and mice of age 6-10 weeks were used. For tumor engraftment, mice were intravenously injected with 5e5 CD19^+^ Raji/ffluc-GFP tumor cells at day -1. At day 7, the mice were treated by intravenous injection of 1e6 primary human CAR T cells generated by CRISPR/Cas9 gene-editing. In-vivo tumor measurement was performed by bioluminescence imaging. Therefore, mice were anesthetized with 2.5% Isoflurane (RAS-4 Rodent Anesthesia system, Perkin Elmer) and 150 mg/kg body weight D-Luciferin-K-Salt (PJK GmbH, Germany) was injected intraperitoneally. 10 minutes post injection, mice were measured with IVIS Lumina Imaging System (PerkinElmer LAS).

Bioluminescence signal analysis was performed by quantification of photons/sec/cm^2^/sr with Living Image 4.5 software (PerkinElmer). Blood analysis was performed for quantification of CAR T cells and CD19^+^ Raji/ffluc-GFP cells in the periphery by flow-cytometry. For blood analysis, erythrocytes were lysed by ACK buffer (Thermo Scientific) in advance. The experiments were approved by the Regierung von Oberbayern (ROB-55.2-2532.Vet_02-17-138).

### Data analysis

Data analysis was performed by Prism 9 Software V9.5.1 (GraphPad), FlowJo software V10.7.1 (TreeStar Inc.), and Living Image 4.5 software (PerkinElmer).

## Author contributions

K. M., L. D., S. P. F., M. E., C. T., V. C., E. B., and M. W. contributed to study design, performed experiments, and analyzed the data. K. S., T. R. M., and D. H. B. provided critical reagents and protocols. C. S., L. G., and M. P. P. designed and directed the project. C.S., L.G., and M. P. P. developed underlying technology. K. M., L. D., S. P. F., M. E., and M.P.P. compiled the manuscript. All authors reviewed this manuscript.

## Data availability

The authors declare that all data generated or analyzed for this study are available within the paper and its supplementary information. Additional raw data are available from the corresponding author upon reasonable request.

## Competing interests

K. M., L. D., S. P. F., M. E., C. T., V. C., E. B., M. W., D. H. B., C. S., L. G., and M. P. P. are currently employed by Juno Therapeutics GmbH, A Bristol-Myers Squibb Company and own stocks of Bristol-Myers Squibb. L. G., C. S., and M. P. P. are listed as inventors on previously filed related patent applications.

