## Supplementary Figure for "Miniaturized CAR knocked onto CD3ε extends TCR function with CAR specificity under control of endogenous TCR signaling cascade"

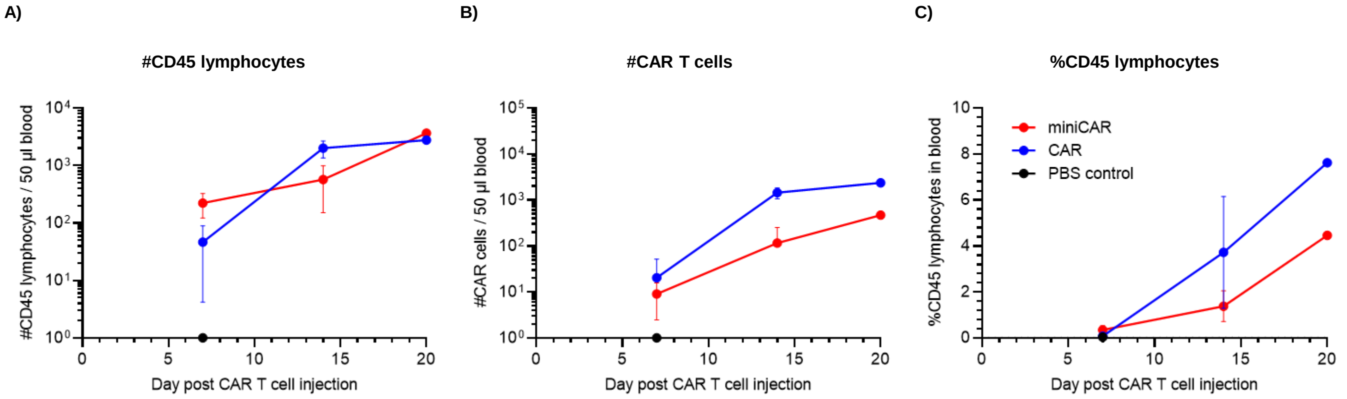


**Supplementary Figure 1. Expansion of CD45 lymphocytes and CAR T cells in tumor-bearing mice. A-C)** NSG-SGM3 mice were engrafted with 5e5 CD19+ Raji/ffluc-GFP cells tumor cells at day -1. At day 7, the mice were treated with 1e6 human primary miniCAR or CAR T cells. The anti-CD19-CAR was introduced by CRISPR-technology respectively. Blood analysis was performed at day 7, 14 and 20 post adoptive T cell transfer. **A)** Flow-cytometric quantification of CD45 lymphocytes in 50 µl blood. **B)** Flow-cytometric quantification of CAR T cells in 50 µl blood. **C)** Proportion of CD45 lymphocytes from total lymphocytes in blood analyzed by flow-cytometry.
